# NeoAtlas and NeoBert: A Database and A Predictive Model for Canonical and Noncanonical Tumor Neoantigens

**DOI:** 10.1101/2025.03.16.643518

**Authors:** Meilong Shi, Qianyi Yan, Wei Zhao, Chuanqi Teng, Fengxian Han, Haobin Chen, Yizhuo Li, Lingyun Xu, Fei Yang, Gang Jin, Yiming Bao, Chunman Zuo, Jing Li

## Abstract

Neoantigens are classified into canonical and noncanonical types. Noncanonical neoantigens include those derived from noncoding regions, transposable elements (TE), and intron retention events, and they have recently gained considerable attention in cancer immunity. We curated 35,574 non-redundant neoantigen-HLA pairs from 14 immunopeptidomes studies, by analyzing unique features and differences across various sources of neoantigens. This knowledge enabled us to develop machine learning models for the prediction of different types of neoantigens. Our data and models are available at a public portal (https://ngdc.cncb.ac.cn/neoatlas) to facilitate broad access and future research. This resource offers advanced functionalities, including integration with epigenome browsers which allow easy navigation of epigenomic datasets to support and confirm the expression of neoantigens. We further demonstrate that combining our database with mass spectrometry analysis can identify noncanonical neoantigens. The resource we constructed holds significant value and promise for the development of neoantigen-based vaccines.

## Introduction

Tumor-specific antigens (TSAs), also referred to as tumor neoantigens, are short peptide antigens displayed by the major histocompatibility complex (MHC) on the surface of tumor cells. These unique TSAs, absent in normal tissues, possess the capacity to elicit T-cell responses specific to tumors, making them promising candidates for cancer vaccines or immunotherapy targets[1].

Canonical TSAs were previously limited to coding gene mutation and fusion[2] derived neoantigens. Consequently, conventional methods for identifying these canonical neoantigens heavily relied on data from whole-genome/whole-exome sequencing[3]. Several extensively curated databases of canonical neoantigens, including CEDAR[4], CAPED (https://caped.icp.ucl.ac.be), TANTIGEN[5], NEPdb[6], dbPepNeo[7], NEOdb[8], and TSNAdb[9], have been broadly explored for their utility in cancer immunotherapy.

Coding regions account for only 1.5% of the human genome. Growing evidence suggests that noncoding genome, as well as noncanonical mechanisms, including transcriptomic variants[10, 11], such as alternative splicing[12, 13], intron retention[14, 15], exitron[14], and transposable element (TE)-derived chimeric transcripts[16, 17] and protein proteosome cryptic peptides (cis/trans proteosome processing[18–20]), contribute to an unknown but potentially large pool of noncanonical neoantigens. However, the collection and curation of noncanonical sources of neoantigens remain limited. For noncoding upstream open reading frame (uORF) derived neoantigens identified through RNA-seq, only one database, SPENCER[21], exists, which collected 2,806 mass spectrometry data from 55 studies. Translation from noncoding RNA (ncRNA) was predicted and segmented into 8-14mer peptides and matched against mass spec data, but there was no evidence supporting their surface presentation. The ProteasomeDB database[22] annotates protein proteasome processing resulted neoantigens. Nevertheless, it entails the *in vitro* synthesis of 80 peptides, followed by proteasome digestion and detection using various mass spectrometry methods. The detection of peptide segments utilizes a method called invitroSPI[22], hence these neoantigens are limited to the synthetic polypeptide substrates.

The detection and sequencing of HLA-bound peptides by liquid chromatography–tandem mass spectrometry (LC–MS/MS) offer a unique advantage to directly identify endogenously processed and presented peptides from cells[23]. Therefore, we decided to focus on analyzing immunoproteomics data from tumor samples and cell lines to accelerate epitope discovery. We collected and curated a large dataset of noncanonical neoantigen peptides encompassing 35,574 non-redundant neoantigen-HLA pairs from different types of RNAs and proteins based on 14 immunopeptidomes studies[10, 11, 14, 15, 18–20, 24–31]. Taking advantage of the largest collection of noncanonical tumor antigens to date, we further compared their antigenic features with those derived from somatic mutations (represented by CEDAR[4]), tumor-associated peptides (represented by HLAthena[32]), and neoantigens originating from infectious diseases (represented by IEDB[33]), including their length, motifs, entropy and hydrophobicity.

We further visited the long-standing challenge of predicting peptide-HLA binding in the context of noncanonical tumor antigen. Current methods for peptide-HLA binding prediction can be categorized into four types: structure-based, scoring function-based, machine learning-based, and integrated methods combining multiple predictors. Structure-based[34] approaches assess the binding structure of HLAs and peptides, while scoring function-based techniques often explore sequence-based attributes[35, 36]. Machine learning-based methodologies extract features from peptides or HLAs and train models for binding prediction. Deep learning methods, including CNNs, RNNs, and attention mechanisms, have all been applied to tackle this problem and they exhibit promising performance[37–48]. Recently, a more advanced transformer-based model was developed to predict peptide–HLA class I binding and to optimize mutated peptides for vaccine design[49]. However, the model was trained only on canonical peptides. We extended this approach and trained a bidirectional transformer model we called NeoBert by integrating both canonical and noncanonical neoantigens, and we obtained promising outcomes.

All datasets and models are accessible through a dedicated website NeoAtlas (https://ngdc.cncb.ac.cn/neoatlas). NeoAtlas is a free, open-source, web-based, database for noncanonical neoantigens and predictive models of antigen-HLA binding. Its primary goal is to aggregate noncanonical neoantigen data to support immunotherapy studies and cancer vaccine development. The website is divided into an RNA section and a protein section to streamline access to noncanonical neoantigens. For RNA-derived neoantigen, we leverage the WashU epigenome browser[50] to highlight the noncoding regions where the neoantigens are derived from. Users can integrate epigenetic annotations from the WashU epigenome browser to examine expression and epigenomic data across tissues and cells, including the ROADMAP data[51]. For neoantigens from protein splicing, we display amino acid sequences from the original proteins and highlight either cis or trans splicing sites. Finally, we have integrated the NeoBert model into the website to assist researchers in predicting the generation and binding affinity of noncanonical antigens.

## Results

### Data sources

Several neoantigen databases have been developed and each has its own unique features and limitations. The classic IEDB database is perhaps one of the first and most comprehensive resource for collecting experimental data on antibody and T cell epitopes in humans and other animals in infectious diseases, allergy, autoimmunity and transplantation[33]. The IEDB also hosts epitope prediction and analysis tools, and has a companion site, CEDAR[4], which houses cancer epitopes. HLAthena is a repository manually curated for tumor-associated antigens eluted from 95 HLA-A, -B, -C and -G mono-allelic cell lines and assayed by mass spectrometry[32]. However, it lacks noncanonical neoantigens, such as intron retentions, TE chimeric, noncoding, and proteasome-spliced neoantigens. Although several databases, such as SPENCER[21] and InvitroSPI[22], address these aspects, they remain underexplored. NeoAtlas closes this gap by collecting a total of 14 cohorts to annotate unique RNA-derived (NeoAtlas-RNA) and protein-derived (NeoAtlas-Protein) neoantigens, resulting in non-redundant 12,349 and 23,282 antigen-HLA binding pairs, respectively (**Fig.1a**).

**Fig. 1.**
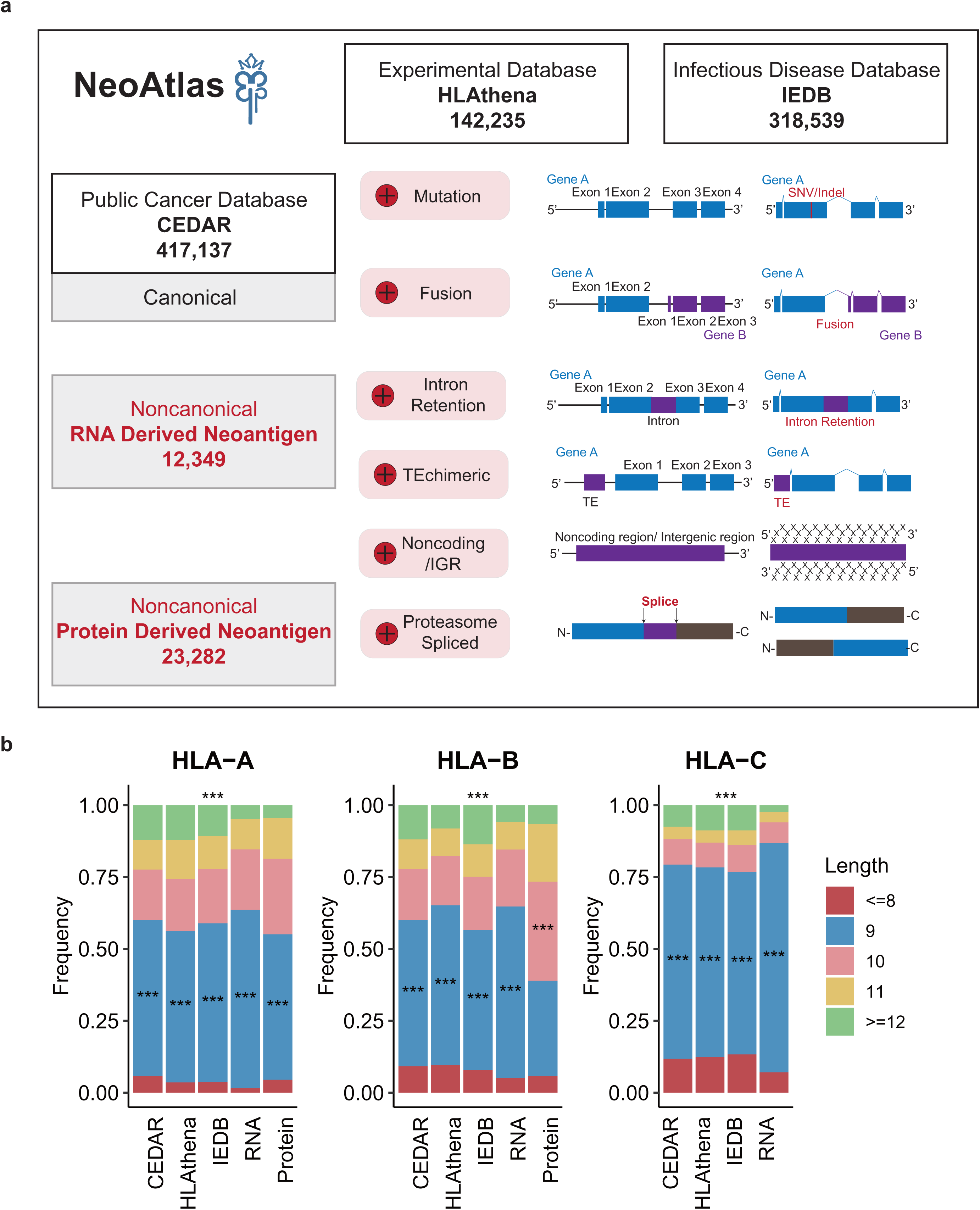
Numbers and length distribution of canonical and noncanonical neoantigens from five databases. a, Three databases representing different types of antigens: IEDB, which collects antigens from infectious diseases; CEDAR, which primarily includes canonical neoantigens from DNA mutations and fusion genes; and HLAthena, which collects tumor antigens from cell lines. The right panel provides a schematic illustration of noncanonical neoantigens derived from RNAs and proteins. **b,** Comparison of peptide length distribution across different databases. The “***” on the top of each HLA indicates that the peptide lengths in the CEDAR, HLAthena, IEDB, NeoAtlas-RNA, and NeoAtlas-Protein databases are all significantly different (chi-square test, p-value < p<2.2e-16). The “***” label for each length indicates that the respective length is dominant within its corresponding database.

RNA-derived neoantigens are primarily detected through RNA-seq combined with mass spec, which identifies noncanonical neoantigens from lincRNAs, 5’ UTRs, TEs, antisense transcripts, and some predicted out-of-frame translated neoantigens in coding regions. In contrast, protein-derived neoantigens are detected using mass spectrometry, which identifies proteasome-derived cis or trans-spliced neoantigens. Cis-proteasome-spliced peptides originate from a single protein, while trans-spliced peptides come from two distinct proteins. Among RNA-derived neoantigens, 25.9% overlapped with coding regions but from alternative coding frames, 39.7% originated from the 5’ UTR, 24.4% from intergenic and noncoding regions (such as exitrons or lincRNA), 5.7% from the 3’ UTR, and 0.6% from transposable elements (TE) (**Supplementary Fig.1a**). Of the NeoAtlas-RNA derived neoantigens, 1,356 were found in IEDB, 7,635 in CEDAR, and 610 in HLAthena. Similarly, 13,611 out of 23,282 protein-derived neoantigens were found in IEDB, 16,357 in CEDAR, and 27 in HLAthena (**Supplementary Fig.1b**). The SPENCER database[21] shared only 35 common neoantigens with NeoAtlas-RNA (**Supplementary Fig.1c**). NeoAtlas-Protein shared only one neoantigen with InvitroSPI[22] (**Supplementary Fig.1d**).

### Variations in peptide lengths, motifs, and extrinsic properties across different databases

The peptide lengths in the CEDAR, HLAthena, IEDB, NeoAtlas-RNA, and NeoAtlas-Protein databases were all significantly different (**Fig.1b**, p<2.2e-16p<2.2e-16, p=2.9e-97). In the HLA-A dataset, the most common peptide length was 9 mer across all databases (p<2.2e-16). In the HLA-B dataset, the Protein database most frequently featured a peptide length of 10 mer (p=0.00236), slightly more than 9 mer, while the other four databases were dominated by 9 mer (p<2.2e-16). In the HLA-C dataset, the most common peptide length across all databases was 9 mer (p<2.2e-16) (**Fig.1b**). The entire dataset spanned 201 HLA subtypes (HLA-A, HLA-B, HLA-C).

Using HLA-A*11:01 as an example among the HLAs with a frequency greater than 20% in the Chinese population, we observed that its most frequent peptide length was 9-mer, but its frequency varied across different databases. In CEDAR, IEDB, and NeoAtlas-RNA, 9-mers accounted for 49%, 47%, and 57% of peptides, respectively. In HLAthena, 9-mer peptides constituted only 39%. Another notable difference was that in NeoAtlas-RNA, peptides of 12-mer or longer made up only 5% of the total, in contrast to 13% in CEDAR, 18% in HLAthena, and 14% in IEDB (**Supplementary** Fig.2, **Supplementary Table 1**). To better understand the basis of differential binding, we compared HLA alleles based on the amino acid preferences of their ligands across different databases (**Supplementary** Fig.3**-6**). For 102 HLA alleles, we analyzed the motifs derived from RNA and protein and compared them to those from the CEDAR, HLAthena, and IEDB databases. The similarity between motifs was measured using Euclidean distance, with a cutoff of 0.236 and a q-value less than 0.05 to determine similar versus dissimilar motifs (see **Methods**). We found that 48 out of 102 (47.1%) HLAs shared peptide motifs across databases. However, 51 RNA-derived peptide-HLA pairs and 12 protein-derived peptide-HLA pairs exhibited significant differences, indicated by their large Euclidean distances (**Supplementary** Fig.7). For example, the 8-mer protein-derived peptides and 9-mer RNA-derived peptides bound to HLA-A68:02 displayed different motifs compared to other databases (**Fig. 2a**). In the case of HLA07:02, the protein-derived neoantigen displayed much reduced preference for P at the second position (**Fig. 2b**). HLA-B27:05 neoantigens from HLAthena showed a preference for F at the last position–(**Fig. 2c**). Similarly, for HLA57:03, the 9-mer RNA-derived peptide showed a stronger preference for W at the last position than peptides from other databases (**Fig. 2d**). To capture the complexity of binding preferences, we calculated the entropy across peptide positions. The analysis revealed that 52 RNA-derived neoantigens and 12 protein-derived neoantigens exhibited differences from other databases, with entropy differences greater than 0.3 (**Supplementary** Figs. 8-9). We also observed significant variations in hydrophobicity among neoantigens across different databases (Welch’s two-sample t-test, two-sided, **Fig. 2e**). In summary, we found that noncanonical neoantigens exhibit differences in binding motif preferences for certain HLAs, which could influence their binding affinity with specific HLA molecules.

**Fig. 2.**
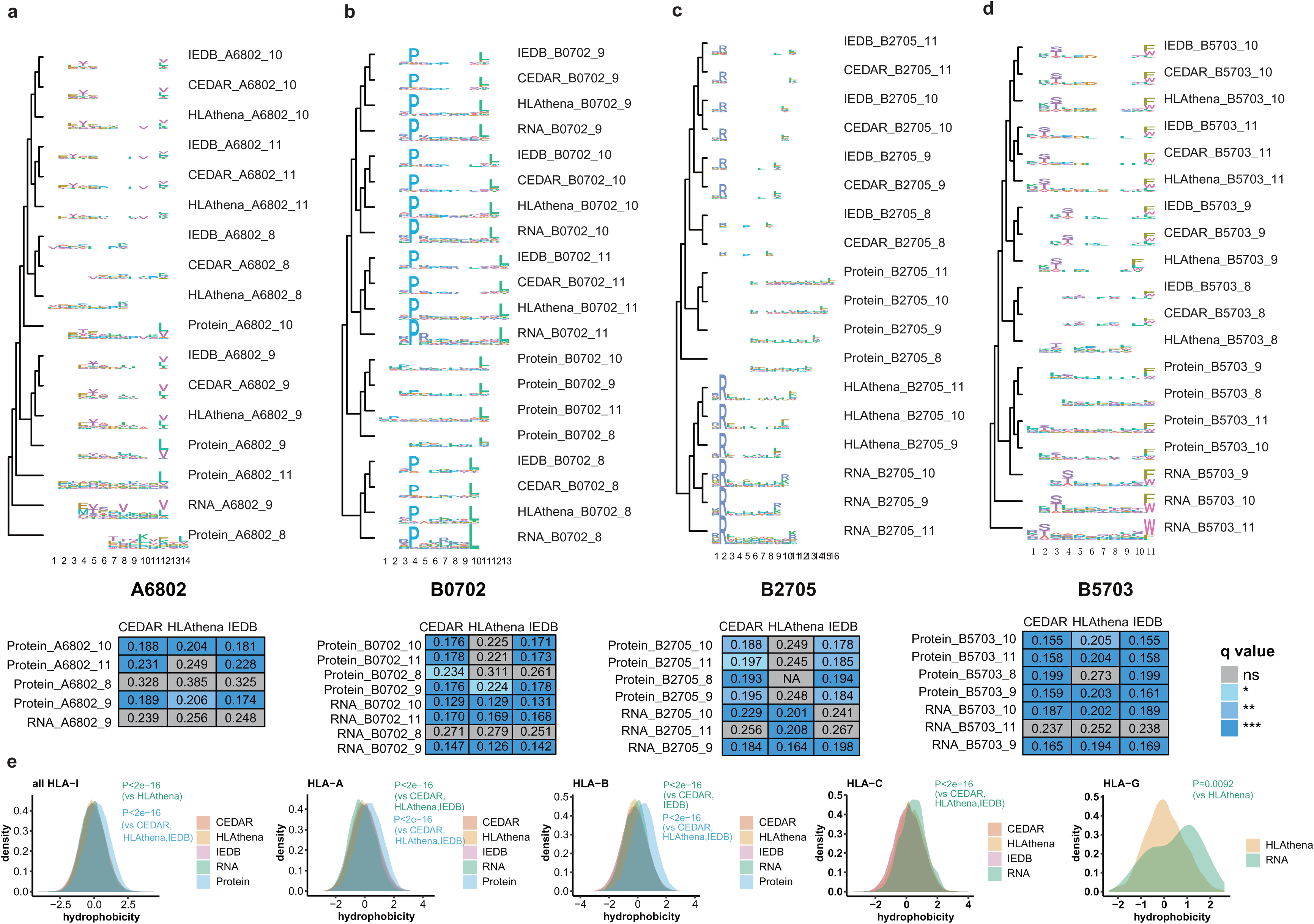
Characterization of canonical and noncanonical neoantigens. a-d, Four examples of amino acid preference comparison of bound peptides of the same HLA across databases. The four representative HLAs are HLA-A68:02, HLA-B07:02, HLA-B27:05, and HLA-C57:03. The amino acid preferences of bound peptides are represented by motif logos. Pairwise comparisons are done by measuring Euclidean distance between the motifs, and the results are clustered using a tree layout. The pair-wise Euclidean distances are included in the table below each tree. Q-values are calculated to determine motif similarity. The *** indicates a q-value of less than 0.001, ** indicates a q-value of less than 0.01, and * indicates a q-value between 0.01 and 0.05. Databases with fewer than 10 peptides were excluded from the analysis. For more details, see **Supplementary** Fig. 3-6**. e,** Distribution of hydrophobicity scores of peptides in each database, stratified by HLA loci (n=204 alleles; 67 HLA-A, 104 HLA-B, 30 HLA-C,3 HLA-G). All comparisons were made using Welch’s two-sample t-test, two-sided.

### Overview of the NeoBert model

Accurately predicting epitope and HLA binding remains a critical and challenging task in immunology, which prompted the development of many tools and models such as NetMHCpan [52]. Very few current models took into consideration of noncanonical antigens, but our work thus far has revealed that distinct features exist between canonical and noncanonical neoantigens. Towards this end, we adopted the Bidirectional Encoder Representations from Transformers (BERT)[53] model to predict the binding affinity of neoantigens and HLA, and named our new model NeoBERT. Our intention was to make NeoBERT applicable to both canonical and noncanonical neoantigens (**Fig.3a**). By treating each HLA-peptide pair as a sentence, NeoBert randomly masks 15% of the amino acids and then predicts these masked amino acids based on the context, to capture bidirectional contextual relations within and between peptide and HLA sequences. Non-linear motif patterns within and between these sequences can be discovered. Further, using a linear binary classifier representing each sentence the positive or negative binding of each HLA-peptide pair can be determined.

**Fig. 3.**
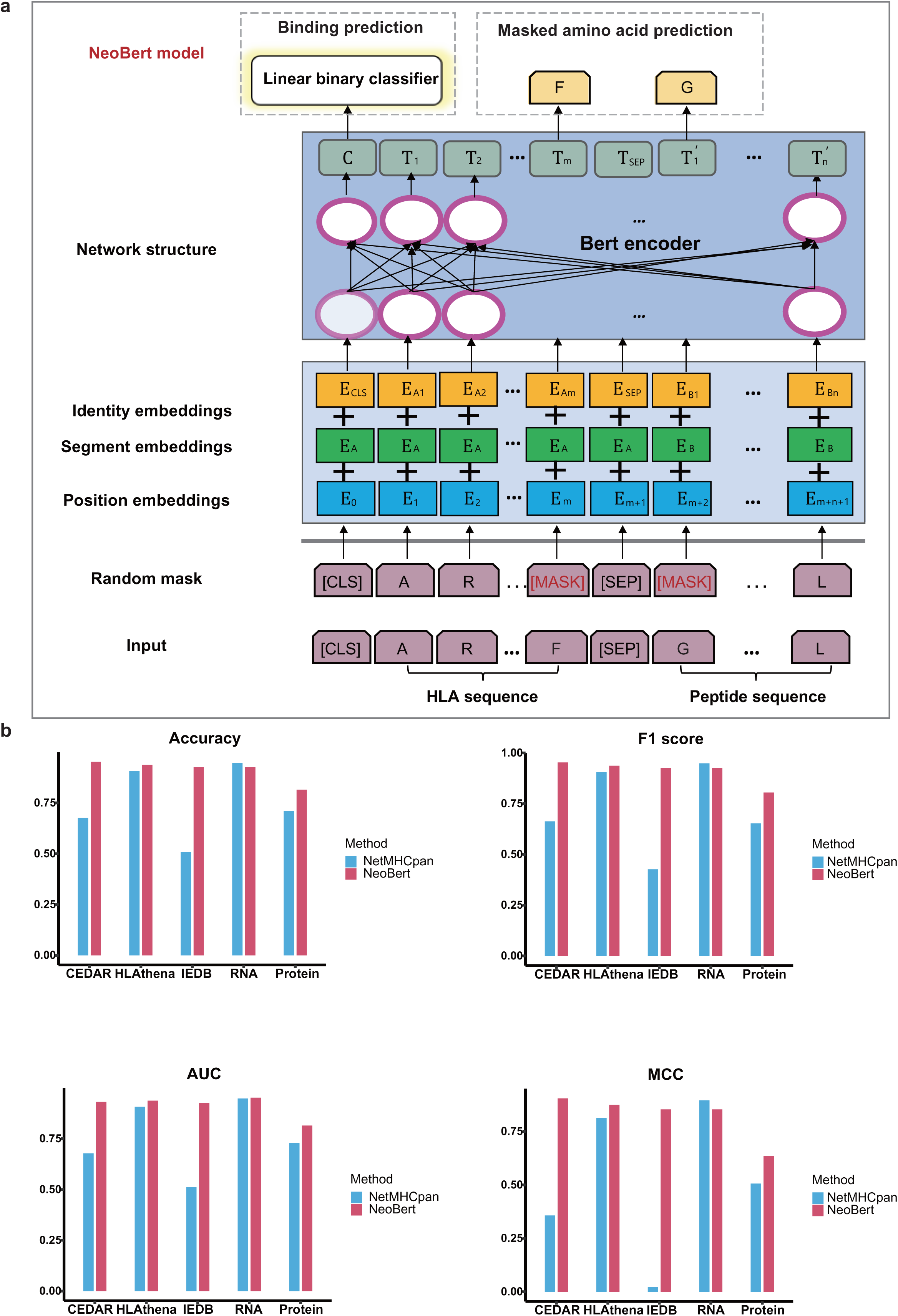
Concept of NeoBert and comparison between the NeoBert model and NetMHCpan. a, The framework of the NeoBert model. Each pair of HLA and peptide sequences is treated as a sentence, with each amino acid represented as a token. The [CLS] token represents the entire sequence, while the [SEP] token separates the HLA and peptide sequences. Random masking is applied to some amino acids in the sequence. For each token, the position embeddings *E_i_*(corresponding to the token’s position in the sequence), segment embeddings *E_A_* (indicating the token’s group), and identity embeddings *E_Ai_* (derived from the amino acid identity using WordPiece embeddings) are summed and fed into BERT to generate token embeddings. The amino acid predictor infers the identities for the masked amino acids, contributing to the loss function. Simultaneously, the [CLS] representation is passed through a linear classifier to predict the binding affinity of neoantigens and HLA sequences. **b,** Bar plots representing the accuracy, F1 score, AUC, and Matthews correlation coefficient (MCC) comparison between NeoBert and NetMHCpan4.1-BA.

After training NeoBert on 70% of HLA-peptide pairs from five different datasets containing canonical and noncanonical neoantigens, we tested its performance against NetMHCpan, a well-established method, on the remaining 30% data. Our benchmarking results (**Fig.3b**) showed that (1) NeoBert achieved an overall accuracy of 93%, compared to NetMHCpan’s 75%; (2) NeoBert outperformed NetMHCpan in predicting both canonical and noncanonical neoantigens, with slightly weaker performance only in RNA-derived neoantigens; and (3) NeoBert provided more consistent prediction results across databases than NetMHCpan did, indicating its capability to capture non-linear diverse motif patterns across different sources.

### Development of NeoAtlas, a noncanonical neoantigen portal

We developed a public data portal, named NeoAtlas, to provide a one-stop shop for investigators to access the noncanonical neoantigen collections as well as our predictive tool, NeoBert **(Fig.4a)**. The current NeoAtlas comprised four core modules: RNA Neoantigens (**Fig.4b**), Cis-Protein Neoantigens (**Fig.4c**), Trans-Protein Neoantigens (**Fig.4d**) and the NeoBert Tool; it encompassed a large collection of data as well as a diverse set of functions to enhance its usability.

**Fig. 4.**
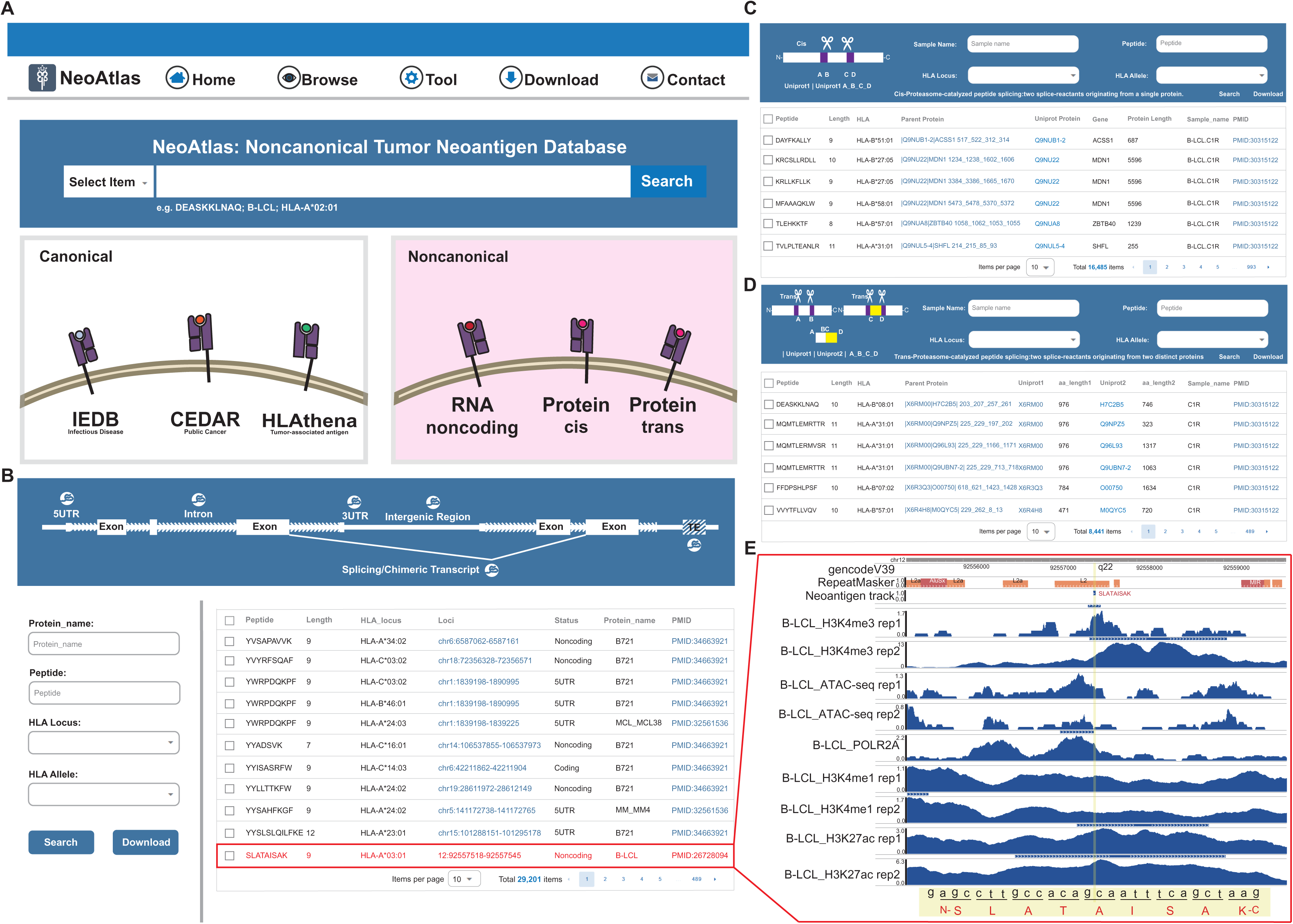
Overiew of the NeoAtlas data portal. a-d, Snapshots of the Home page, NeoAtlas-RNA, NeoAtlas cis, and NeoAtlas trans protein pages. **e,** Example of the NeoAtlas-RNA neoantigen “SLATAISAK” location and integrated epigenome data from Cistrome DB displayed in the WashU epigenome browser. The tracks shown include ATAC-seq, H3K4me3, H3K4me1, and H3K27ac ChIP-seq data, along with POLR2A ChIP-seq of B-LCL. And a default neoantigen track is also displayed.

NeoAtlas supports various types of searches including neoantigen types and peptide sequences, and supports data downloads. The RNA Neoantigens module of NeoAtlas contained 33,725 RNA neoantigens including their peptide sequences, sequence lengths, genomic loci, overlapping genomic features, cell lines, and other details. Users may use a series of filtering criteria such as protein name, peptide sequence, HLA locus, and HLA allele to explore, access and download information related to the neoantigen of interest. PMID links were provided to let users navigate to the original data source and literature. The Cis and Trans Protein Neoantigens module of NeoAtlas contained 9,928 and 4,889 entries respectively. These entries featured peptide sequences, sequence lengths, parent proteins, protein lengths, genes, cell lines, and other relevant information. The NeoBert Tool module provided online and real-time analytical tools for predicting the binding affinity of noncanonical antigens. Upon entering peptide sequences in FASTA format and selecting information about alleles, and lengths, NeoBert would return predicted results for the neoantigen instantaneously.

NeoAtlas created a range of user-friendly web interfaces for easy visualization of noncanonical tumor antigens. For RNA-derived neoantigens, the “loci” link enabled visualization of peptide genomic locations via the WashU Epigenome Browser. Users can take advantage of the rich public epigenomic data to annotate the RNA from which the predicted noncanonical neoantigen derived. For example, in Fig 4e we illustrated the peptide “SLATAISAK” derived from a noncoding transcript from B-Lymphoblastoid cell lines (B-LCL). This included promoter signals (H3K4me3 and H3K27ac) and POLR2A ChIP-seq data, providing strong epigenetic support of transcription from these noncanonical starting sites (**Fig.4e**). For each protein-derived neoantigens, NeoAtlas included a hyperlink to showcase the nucleotide sequence of the gene and highlight the peptide segment.

### Using NeoAtlas to assist mass spectrometry-based neoantigen detection

Mass spectrometry-based neoantigen identification has gained a lot of tractions in the field, but challenges remain. A typical strategy relies on database searches in which theoretical spectra from protein sequences in the database were generated and compared with experimental spectra to find matches. Therefore, a comprehensive and accurate database is crucial for successful mass spectrometry-based neoantigen identification. To test the effectiveness of NeoAtlas, we generated MHC-I pull-down mass spectrometry data using CCLE cell line AsPC-1 and compared it with different neoantigen databases (IEDB, CEDAR, HLAthena, NeoAtlas-RNA, NeoAtlas-Protein). After excluding 3,984 peptides mapped to benign antigen database[54] and 4,062 peptides mapped to Uniprot database, we identified 902 HLA pairs matched to IEDB, 1,153 to CEDAR, 554 to HLAthena, 54 to NeoAtlas-RNA, and 41 to NeoAtlas-Protein (**Fig.5a, Supplementary Table 2**). Using AsPC-1 RNA-seq data we conducted *de novo* transcript assembly (**Fig.5b**) and validated three NeoAtlas-RNA entries with RNA-seq data: SVAPASGAF&HLA-B*15:01 (**Fig.5c**), LVTDLDTEY&HLA-B*15:01 (**Fig.5d**), and VAAESHPL&HLA-C*03:04 (**Fig.5e**).

**Fig. 5.**
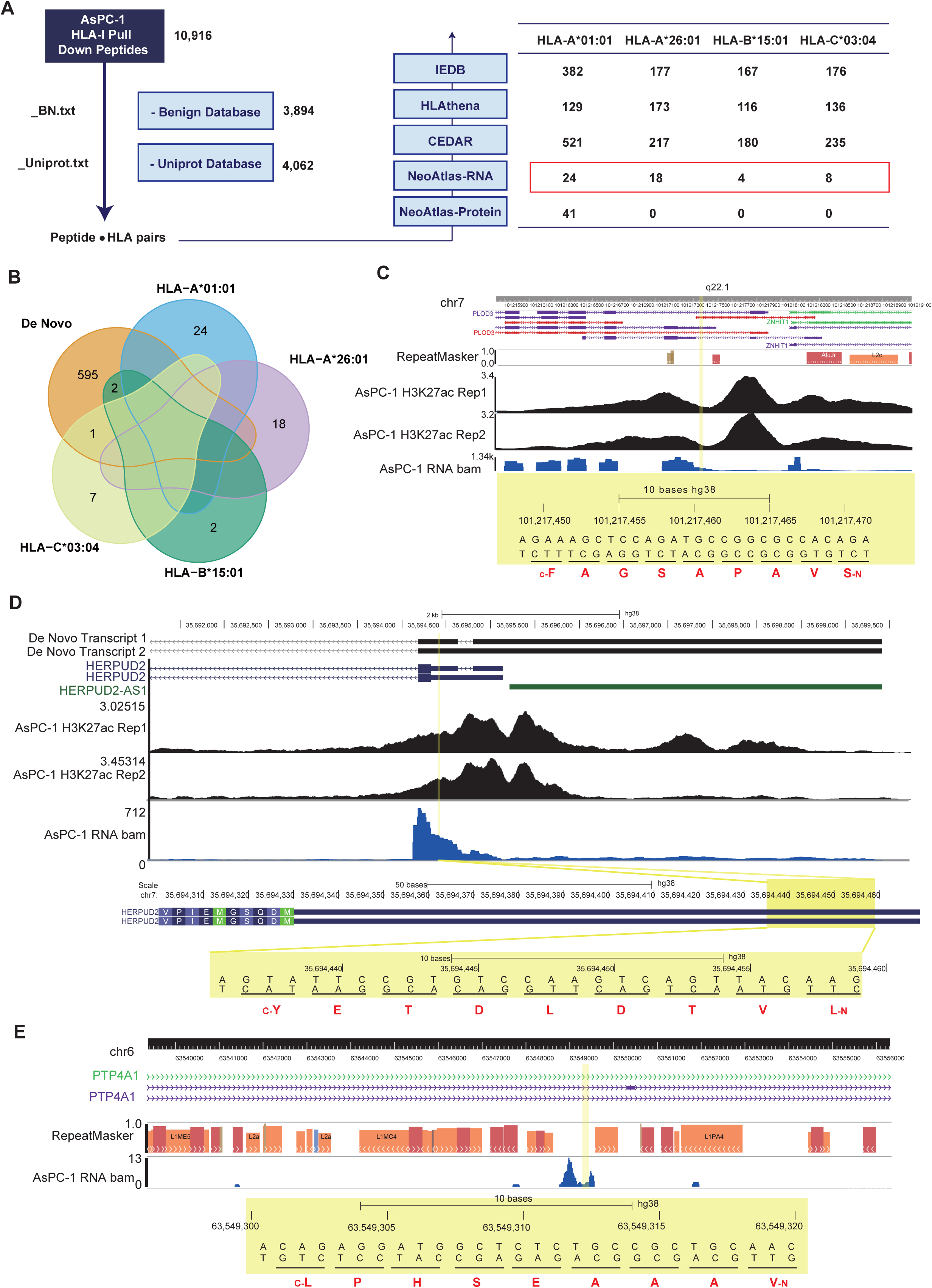
Immunopeptidomics data of AsPC-1 and their analysis using five databases. a, The pipeline for HLA-I pull-down peptide identification. Peptides found in benign and UniProt databases were filtered out, leaving matched peptides and HLA pairs. The overlap numbers across the databases are listed in the right panel. **b,** Venn diagram of peptide-HLA pairs compared with *de novo* assembled transcripts from AsPC-1 RNA-seq data. **c,** The peptide SVAPASGAF derived from AsPC-1 RNA-seq reads, supported by data shown in the WashU Epigenome Browser. The chromosome location, gene annotation, two AsPC-1 H3K27ac ChIP-seq datasets from Cistrome DB, and CCLE AsPC-1 RNA-seq data are displayed. The coding region of the peptide is highlighted in yellow, with “N” representing the N-terminal and “C” representing the C-terminal of the peptide. **d,** The peptide LVTDLDTEY derived from AsPC-1 RNA-seq reads, supported by data shown in the WashU Epigenome Browser. The chromosome location, gene annotation, two AsPC-1 H3K27ac ChIP-seq datasets from Cistrome DB, and CCLE AsPC-1 RNA-seq data are displayed. The coding region of the peptide is highlighted in yellow, with “N” representing the N-terminal and “C” representing the C-terminal of the peptide. **e,** The peptide LVTDLDTEY derived from AsPC-1 RNA-seq reads, supported by data shown in the WashU Epigenome Browser. The chromosome location, gene annotation and CCLE-AsPC-1 RNA-seq data are displayed. The coding region of the peptide is highlighted in yellow, with “N” representing the N-terminal and “C” representing the C-terminal of the peptide.

The three peptides (SVAPASGAF, LVTDLDTEY, and VAAESHPL) were also present in our NeoAtlas database (**Supplementary** Fig. 10). For example, VAAESHPL (**Supplementary** Fig. 10a) was found to bind with 11 HLA alleles[10, 11, 30], and similarly, LVTDLDTEY (**Supplementary** Fig. 10c) has been discovered in melanoma, GBM, rectal cancers, and the B721 cell line[10, 30]. SVAPASGAF[10, 30] (**Supplementary** Fig. 10b) has also been previously reported in two studies, covering B721, GBM, and melanoma samples.

## DISCUSSION

Recently, mass-spectrometry-based immunopeptidomics has facilitated the discovery of noncanonical antigens—those derived from sequences outside protein-coding regions or generated by noncanonical antigen-processing mechanisms. When combined with transcriptomics and ribosome profiling, this approach enables the identification of thousands of noncanonical peptides, many of which may be exclusively detected in tumors.

For example, proteasome-derived peptides show a clear preference for specific lengths of intervening sequences (the sequences excised between two splice reactants). Additionally, the lengths of the N- and C-terminal splice reactants are almost equally distributed, with a notable preference for a length of two residues in the N-terminal splice reactant, particularly for 9-, 10-, and 11-mer peptides[22]. Since the second residue of the antigenic peptide often corresponds to the anchor site of specific HLA-I molecules, it is speculated that a preference for specific amino acids during ligation could exert evolutionary pressure on HLA allotype selection[55]. This hypothesis aligns with previous theories regarding proteasome-dependent peptide hydrolysis and the C-terminus of HLA-I–restricted peptides[19].

Given these distinct protein processing mechanisms, noncanonical neoantigens likely exhibit unique features compared to canonical neoantigens. Therefore, the primary goal of this study was to collect and analyze the features of noncanonical neoantigens. We aimed to determine how they differ from canonical neoantigens. Understanding these differences could provide an opportunity to develop new binding affinity prediction tools. Furthermore, visualizing our neoantigen data across the entire genome would provide a convenient platform for investigators and facilitate deepening of our knowledge of noncanonical peptides.

Our study has some limitations. First, in terms of data processing, we only collected the final processed data from existing literature in order to maintain high confidence in our neoantigen collection, instead of downloading and reprocessing raw RNA-seq or Ribo-seq data from each study and integrating them into our database or genome browser. Additionally, HLA pull-down data were limited by not specifying which HLA binds to the neoantigen, thus there was still a disconnection between the neoantigen and specific MHC molecule pairs.

A second limitation is related to data content, where the majority of the peptides we collected were MHC I peptides, and MHC II peptides only made up of a smaller percentage. We hope to collect more MHC II peptides in the future. Furthermore, we did not include some minor groups of noncanonical neoantigens, such as circular RNA and bacteria-derived neoantigens. We hope to explore these areas more thoroughly in the future.

Finally, understanding how TCR interacts with noncanonical neoantigen-MHC complexes remains a significant gap in the field. More research and studies are needed to explore this area and to advance TCR therapeutic development.

In conclusion, we hope the NeoAtlas and NeoBert resource will help shed light on the noncanonical aspects of neoantigens and contribute to future clinical applications.

## Materials and methods

### Collection of datasets

#### For RNA-derived neoantigens data collection

We collected data from 10 previous literatures[10, 11, 15, 24, 25, 27, 28, 30, 31], annotating the following information for each entry: “PMID”, “Cell.line”, “Peptide”, “Length”, “Genome”, “Chromosome”, “Start”, “Stop”, “Loci”, “Transcript_strand”, “Overlapping genomic feature”, “HLA_locus”, “HLA_allele”, “Ensembl_gene_id”, and “Protein_name”. From each paper, we extracted these specified information, considering special circumstances as follows:

For “PMID:31745090”, additional information for “Cell.line”, “ Overlapping genomic feature “, “HLA_allele”, and “Protein_name” for some peptides was supplemented from “41467_2019_13035_MOESM9_ESM.xlsx”.

For “PMID:30114007” and “PMID:33861991”, we downloaded the current version of the UniProt annotation file from https://www.uniprot.org/ and added the corresponding “Ensembl_gene_id” based on “geneName”.

For “PMID:32157095”, data was categorized into HLAp.category==”TE” and HLAp.category!=”TE”. For TE peptides, those starting with “sp” in “Transposable_Element” were directly searched in UniProt to obtain the corresponding Gene, transcript, and Loci. For “Transposable_Element” peptides not starting with “sp”, Gene_name, Loci, and Strand were extracted from “Transposable_Element”, and the Loci were lifted over to hg38. For non-TE peptides, the corresponding Loci were matched from gencode.v24lift37.annotation.gtf (downloaded from https://www.gencodegenes.org/human/release_24lift37.html).

For “PMID:34663921”, “Mapped protein” and “ORF_ID” were matched by merging “41587_2021_1021_MOESM5_ESM.xlsx” and “41587_2021_1021_MOESM6_ESM.xlsx”. Only B721 detected results were retained after merging, which were then combined with Cancers+MHC-I+peptides(1).xlsx to obtain peptide information. Annotation files were downloaded from the NCBI Gene Expression Omnibus (GEO) GSE143263, specifically GSE143263_nuORFdb_v1.0_annotations.xlsx, to annotate peptides.

Additionally, all chromosome loci were liftover to hg38 (PMID: 21728094, 32047025, 32157095, 31460334). Furthermore, genomic feature annotations were standardized. The data from the 10 studies were then integrated. If a peptide had multiple location annotations, the information was split into multiple rows.

### Protein derived neoantigen

We collected data from 4 previous studies[18–20, 26]. We downloaded annotation data from UniProtKB/Swiss-Prot (https://www.uniprot.org/) to annotate proteins and obtain their corresponding genomic coordinates. For the two cis datasets, “PMID:27846572” and “PMID:30315122”, we standardized the Parent protein(s) format to “Entry|Entry.name location” and added protein annotations under “GeneUniprot”.

For “PMID:32938616”, we extracted peptide and Protein name. For “PMID_30346791”, the “Full Sequence” and “Protein Name” columns sometimes contained multiple peptides and proteins connected by “or”. We split these entries to list each Sequence and Protein Name relationship individually, then matched them with protein, Gene name, and Ensemble from the UniProt annotation file. For unmatched entries, we downloaded the corresponding protein’s FASTA file from UniProtKB. If still unmatched, we downloaded the FASTA file from Unisave. The download commands used were: “wget https://rest.uniprot.org/uniprotkb/$i.fasta” and “wget https://rest.uniprot.org/unisave/$i?format=txt”. ($i can be replaced with the corresponding protein name).

### IEDB, CEDAR, HLAthena data access and preparation

The public data were cleaned as previously described[32]. A curated set of previously identified HLA class I ligands was downloaded from IEDB at http://www.iedb.org/downloader.php?file_name=doc/mhc_ligand_full**.zip (accessed on May 2024). The HLAthena were downloaded at http://350HLAthena.tools and CEDAR data were downloaded at https://cedar.iedb.org/database_export_v3.php. Records were filtered to MHC allele class = I, Epitope Object Type = Linear peptide and Allele Name consistent with human HLA class I nomenclature with four-digit typing (that is, regex: “^HLA-[ABCG]\\*[0–9]{2}:[0–9]{2}$”). Peptides with quantitative measurements in units other than nM were removed and so were the following three assay types due to inconsistency between predicted and actual affinity: ‘purified MHC/direct/radioactivity/dissociation constant KD’, ‘purified MHC/direct/fluorescence/half maximal effective concentration (EC50)’ and ‘cellular MHC/direct/fluorescence/ half maximal effective concentration (EC50)’. A peptide was considered a binder if it had a quantitative affinity of <500 nM or qualitative label of ‘Positive’, ‘Positive-High’, ‘Positive-Intermediate’ or ‘Positive-Low’. In cases where multiple records were available for the same {peptide, allele} pair, we either took the mean affinity or removed the peptide when the difference between the maximum and minimum log-transformed affinities (1 − log(nM)/log(50,000)) was >0.2. Similarly, peptides with multiple qualitative records were removed if the same number of positive and negative labels were found or kept otherwise.

### Entropy

The entropy was calculated as previously described[32]. The entropy at each peptide position (i.e.1 through n for n-mer peptides,n∈{8,9,10,11}) was calculated for each allele based on all peptides identified. The computation was performed with MolecularEntropy() function from HDMD R package. For each HLA, we determined the absolute difference in entropy between motifs of varying lengths at each position in CEDAR, HLAthena, IEDB, RNA, and Protein datasets. We then analyzed the position with the maximum entropy difference (the value on the y-axis in the figure represents the entropy at this position). An absolute difference in entropy greater than 0.3 was considered indicative of significant motif differences between the databases.

### Peptide motif distance

We utilized Euclidean distance (EUCL) to quantitatively assess motif-motif similarity, as referenced by Choi et al.[56]. To compare the similarity of motifs across different databases, we employed the Tomtom algorithm[57], using EUCL as the basis for comparison. Specifically, we created a distance matrix of motifs from different databases using the compare_motifs function from the R package universalmotif (method set to “EUCL”). We then generated and visualized a phylogenetic tree of motifs based on the distance matrix using the motif_tree function from the same package (layout set to “rectangular”). Additionally, we compared the similarity of motifs between different databases using the runTomTom function from the R package memes (with the parameter dist set to “ed”). We selected a EUCL cutoff of 0.236 because it corresponds to the 75th percentile of all distances, and a q-value of less than 0.05 to distinguish between similar and dissimilar motifs. Through these methods, we effectively quantified and visualized the similarity of motifs across different databases.

### NeoBert model

The core principle of NeoBert is to capture the contextual information within and between peptide-HLA sequences using bidirectional encoder representations from transformers. NeoBert consists of four sequential components **(Fig.3)**: (i) each pair of HLA and peptide sequences is combined into a single sentence, with randomly masked amino acids used as input; (ii) embedding block adds positional embedding to the identity (i.e., amino-acid) embeddings and segment embeddings to generate the sequence embeddings; (iii) encoder block comprises 12 BertLayers, each containing masked multi-head self-attention mechanisms and a feature optimization block to learn representations; and (iv) a linear binary classifier predicts the binding probability using the aggregated sequence representation, while an amino-acid predictor model infers masked amino acids based on the representations of each corresponding position. Specifically,

***(i) Input sequence***

We represented each pair of HLA and peptide sequences as a sentence, i.e., *X*= ([*CLS*], *HLA sequence*, [*SEP*],*peptide sequence*). The final hidden feature of the first token ([CLS]) serves as the sequence representation for classification, while the ([SEP]) token differentiates the two sequences. Randomly masked amino acids for both sequences were used to train a deep bidirectional representation. The input peptide and HLA sequences were padded with zero to a length of 128 to accommodate variable input lengths.

***(ii) Sequence embedding***

For each token, we added learned embeddings (*E_A_* or *E_B_* ϵ *R*^H^) to indicate whether it belongs to HLA or peptide, facilitating the segmentation of these two sequences, where *H* is the dimension of embeddings. We used WordPiece embeddings [58] to encode the identity of amino-acid (*E_Ai_* or *E_Bj_* ϵ *R*^H^). Additionally, positional embeddings (E_i_ ϵ *R^H^*) were incorporated to encode the position of amino acid in the whole sentence (*X*). Hence, the input representation of the *ith* token in *X* (or *mth* token in HLA) is created by summing its token, segment, and position embeddings as follows:

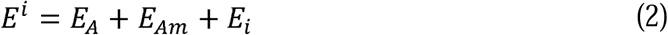

***(ii) Encoder block***

We leveraged the multi-head self-attention technique within a concatenated sequence to effectively incorporate bidirectional cross-attention between the two sequences.

The mechanism involves mapping the query *Q* to a set of key-value (*K*- *v*) pairs and obtaining an output, with *K*-*v* pairs storing sequence elements in memory. The attention score is based on the correlation or similarity between *Q* and *K*, signifying the importance of information. A higher attention score (i.e., *v*) indicates a stronger focus on the corresponding information. Furthermore, the feature optimization block combines fully connected layers ([768, 3072, 768]) with

LayerNorm and Dropout to learn enhanced representations (*H*^i^) for the *ith* token. For concatenated sequences of HLA and peptide with a length less than 128, we padded them with zeros to 128. Hence, non-amino-acid characters do not play a role in calculating the attention.

***(iv) Prediction block***

We adopted two tasks to fine-tune our NeoBert model: HLA-peptide binding prediction and masked amino acids prediction. Specifically, we fed the aggregated sequence representation of each HLA-peptide pair into a sigmoid layer to predict the label:

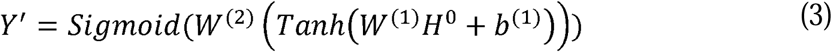

where *w*^(l)^, *w*^(2)^, and *b*^(l)^ are learnable parameters, and *H*^O^ indicates the representations of the special token [CLS]. The corresponding loss function is outlined as follows:

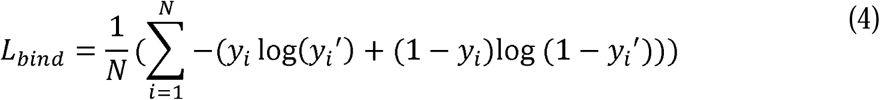

where *N* denotes the set of HLA-peptide pairs containing positive binding data and negative data generated similarly to previous studies[49]. Simultaneously, a masked amino acids model predicts the probabilities of 20 different amino-acids at the *ith* position of the sequence based on its representations:

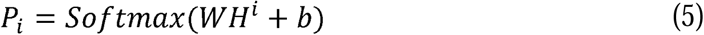

where *w* and *b* are learnable parameters, and *H*^i^ represents the representation of the *ith* token. Its loss function is summarized as follows:

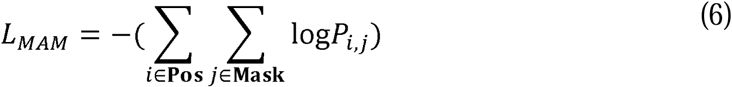

where **Mask** indicates the set of positions of all masked amino-acids, and **Pos** represents the set of positive binding pairs. Hence, the total loss function for NeoBert model is summarized as follows:

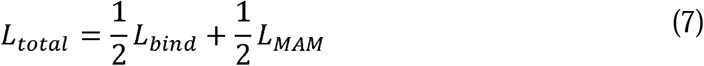

### NetMHCpan prediction

We collected peptide sequences (8-11mer) and their corresponding HLA-I alleles’ data from the database. Using NetMHCpan-4.1, we predicted the binding affinity of each peptide to its corresponding HLA allele, adjusting the peptide length parameter accordingly. For each peptide, the best, i.e., the lowest, percentile rank value was retained as the prediction result. A percentile rank cutoff of 2 was used for weak binders and 0.5 for strong binders.

### NeoAtlas website

NeoAtlas was constructed using Spring Boot (https://spring.io/projects/spring-boot) and deployed in the Centos Linux environment. The backend database resides in MySQL (https://www.mysql.com/). The frontend web interfaces were crafted with Vue.js (https://vuejs.org/).

### Metrics evaluation

We evaluated the performance of each predictive model based on the following metrics: accuracy, Matthews correlation coefficient (MCC), *F*_l_ score, and the area under the curve (AUC) of the receiver operative characteristics curve. Specifically,

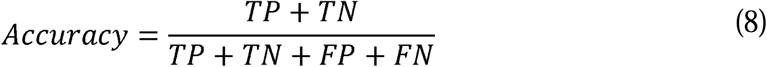

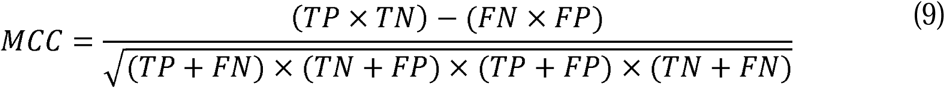

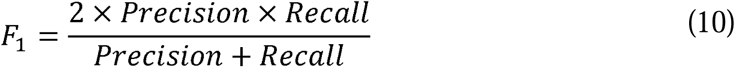

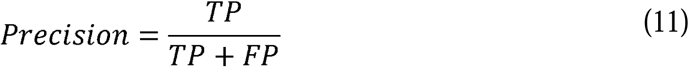

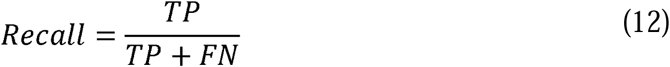

where TP, FP, FN, TN represent true positives, false positives, false negatives, and true negatives.

### Cell line and cell culture

The AsPC-1 cell line was obtained from the National Collection of Authenticated Cell Cultures. AsPC-1 was cultured in RPMI 1640 (Gibco, 11875119) supplemented with 10% fetal bovine serum (Gibco, 10099-141) and maintained in a humidified atmosphere containing 5% CO_2_ at 37°C.

### Immunopeptidomics

The eluted peptides were delivered to timsTOF Pro2 (Bruker Daltonik, Germany), and their MS spectra were acquired in the DDA mode. One MS1 survey TIMS-MS and ten PASEF MS/MS scans were acquired per acquisition cycle. The parameter settings in the machine were adjusted for the best performance. Ion accumulation and ramp time in the dual TIMS analyzer were both set to 100 ms, and we analyzed the ion mobility range from 1/K0 = 0.75 Vs cm−2 to 1.30 Vs cm^−2^. Precursor ions for MS/MS analysis were in a total m/z range of 100–1700 by synchronizing quadrupole switching events with the precursor elution profile from the TIMS device. The ion fragmentation mode was CID and the dynamic exclusion time was set to 30s.

### HLA class I antigen isolation from AsPC-1

We followed a published protocol with a couple of modifications to generate HLA class I antigen pulldown samples[59]. Approximately 2 × 10^7 AsPC-1 cells were used. Subsequently, we used an HLA-I immunopeptide enrichment kit to complete the enrichment and elution of MHC-peptide complexes. The cell samples were added to lysis buffer and incubated on ice for 60 minutes, followed by 5 minutes of sonication. After 21,000g at 4LJ°C centrifugation, the supernatant was collected for subsequent enrichment of immunopeptides. Peptide samples dissolved in formic acid water were then separated using a Bruker Nano-Elute high-performance liquid chromatography system. Mass spectrometry data acquisition was performed using a Bruker timsTOF Pro2 high-resolution mass spectrometer in DDA mode. The data acquisition duration was 60 minutes, with a mass-to-charge ratio scanning range of 100-17000 m/z. PASEF settings included 10 MS/MS scans (total cycle time 2.22 seconds) and an ion intensity threshold of 2500. Raw data in the form of mass spectra were generated.

### HLA mass spectrometry analysis (Blast pipeline)

The mass spectrometry second-level spectral data were acquired using the DDA acquisition mode. All raw data files were processed to obtain peptide identification results. Peptides with lengths ranging from 7 to 16 amino acids were selected, with a delta retention time (deltaRT) range of [-19, 13] and an MS2 ion intensity threshold greater than 0.6. Subsequently, the identified peptides were compared against the Human Benign Proteome database (PXD03878), followed by the Human Proteome database (available at https://www.uniprot.org/) and database we curated (IEDB, CEDAR, HLAthena and NeoAtlas).

### Conservation analysis of the HLA class I antigen and visualization with the WebLogo

We inputted the peptides obtained from the previous comparisons into WebLogo[60] according to their lengths to visualize peptide motifs.

### CistromDB data selection and integration with WashU epigenome browser

ChIP-seq data were obtained from Cistrome Data Browser[61] available at http://cistrome.org/db/<u>#/</u>. ChIP-seq data[62] were visualized by using WashU epigenome browser[63] available at https://epigenomegateway.wustl.edu. Cistrome DB IDs of analyzed ChIP-seq data are 3756, 59349, 78766, 80802, 85770, 89337, 92412, 92226 and 92419.

### AsPC-1 RNA-seq processing and *de novo* assembly

The raw transcriptomic data of AsPC-1 were obtained from the Cancer Cell Line Encyclopedia (CCLE)[64] (https://sites.broadinstitute.org/ccle/datasets, SRR8615412). Quality control was conducted using fastp v0.23.2[65]. The cleaned reads were aligned using STAR v2.7.7a[66] with a two-pass alignment strategy based on the GRCh38 reference genome and GENCODE v42[67] annotation. Novel junctions were merged to create an updated reference for the second pass of the STAR alignment. StringTie v1.3.3b[68] was then used to perform *de novo* assembly based on the STAR two-pass alignment results.

## Acknowledgements

This study was supported by the “National Key R&D Program of China” fund (2022YFA1305700, J. L.) granted by the Chinese Ministry of Science and Technology, the National Natural Science Foundation of China (No. 32300523, C.Z.), Shanghai Sailing Program (No. 22YF1401700, C.Z.), Fundamental Research Funds for the Central Universities (No. 2232022D-30, C.Z.). We acknowledge Daofeng Li for his assistance in linking the WashU Genome Browser.

## Author contributions

**Meilong Shi:** Data Collection, Statistical analysis, Writing - Original Draft, Visualization. **Qianyi Yan:** Methodology, Writing - Original Draft. **Wei Zhao:** Software, Visualization. **Chuanqi Teng:** Data Collection**. Fengxian Han:** Data Collection, Statistical analysis. **Haobin Chen:** Methodology. **Yizhuo Li:** Data Collection. **Lingyun Xu:** Software, Data Collection. **Fei Yang:** Project administration. **Gang Jin:** Writing – review & editing, Supervision. **Yiming Bao:** Writing – review & editing, Supervision. **Chunman Zuo:** Writing – review & editing, Supervision, Funding acquisition. **Jing Li:** Conceptualization, Writing - Review & Editing, Supervision, Project administration, Funding acquisition. All the authors have read and approved the final manuscript.

## Additional information

**Supplementary information** is available for this paper.

**Correspondence and requests for materials** should be addressed to J.L

**Supplementary Fig. 1 | Genomic features of neoantigens in NeoAtlas and their overlap with other databases. a,** Pie charts illustrating the genomic distribution of neoantigens in NeoAtlas-RNA. **b,** Venn diagram showing the overlap of peptides across five different databases. **c,** Venn diagram depicting the overlap between NeoAtlas-RNA and the SPENCER database. **d,** Venn diagram displaying the overlap between NeoAtlas-protein and the InvitroSPI database.

**Supplementary Fig. 2 | Length distribution of peptides in five databases.** The length distribution and overlap comparison between different databases are shown (n=201 alleles; 67 HLA-A, 104 HLA-B, 30 HLA-C). The bottom heatmaps display the relative median population frequencies per allele across populations worldwide. See also Supplementary Table 1.

**Supplementary Fig. 3 | Logo plots for 8-mer peptides in five databases.**

**Supplementary Fig. 4 | Logo plots for 9-mer peptides in five databases.**

**Supplementary Fig. 5 | Logo plots for 10-mer peptides in five databases.**

**Supplementary Fig. 6 | Logo plots for 11-mer peptides in five databases.**

**Supplementary Fig. 7 | Motif clustering and similarity calculation for peptides across five databases.**

**Supplementary Fig. 8 | Entropy difference between RNA-derived neoantigens and those from three other databases.**

**Supplementary Fig. 9 | Entropy difference between protein-derived neoantigens and those from three other databases.**

**Supplementary Fig. 10 | Identification and validation of neoantigen peptides in AsPC1 cells using NeoAtlas database. Three peptides discovered in our AsPC1 cell line were also identified in the NeoAtlas database. The corresponding figure provides detailed information on the relevant studies’ PubMed IDs, sample sources, protein origins, and associated HLA alleles and loci.**

**Supplementary Table 1 | Frequency distribution of peptide lengths and HLA distribution across different human populations.** This table provides a comprehensive analysis related to Fig.1b and Supplementary Figure 2, detailing the frequency distribution of peptide lengths. Additionally, it includes HLA distribution among various human races.

**Supplementary Table 2 | Identification and annotation of peptides from AsPC-1 immunopeptidomics.** This table presents the peptides identified through HLA-I pull-down immunopeptidomics of AsPC-1 cells. It also shows the overlap of these peptides with the databases, in various combinations, alongside the HLA types of AsPC-1 cells predicted from RNA-seq data.

