## Supplementary figures and images for "NeoAtlas and NeoBert: A Database and A Predictive Model for Canonical and Noncanonical Tumor Neoantigens"

### Supplementary Figure 1

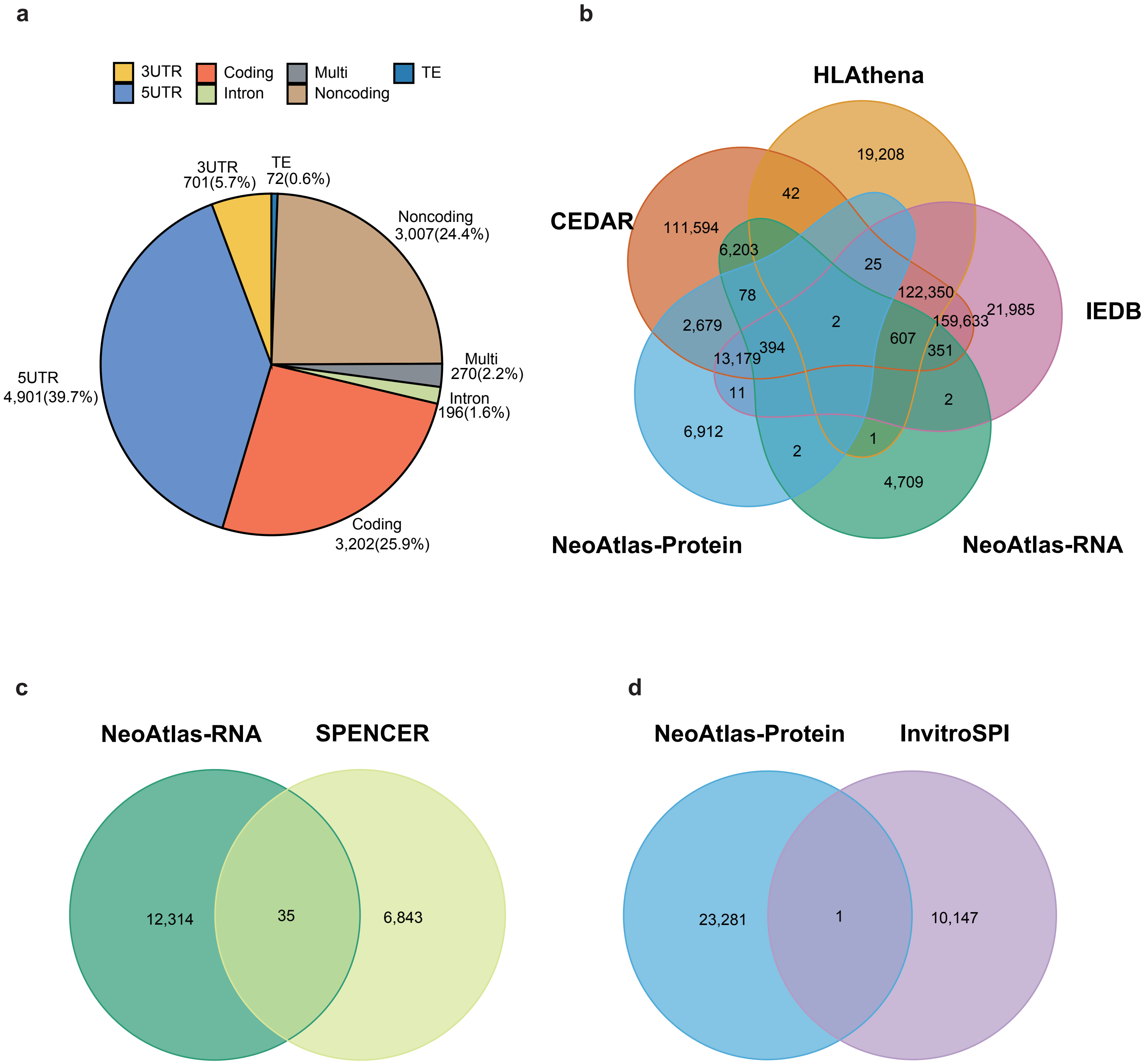

### Supplementary Figure 2

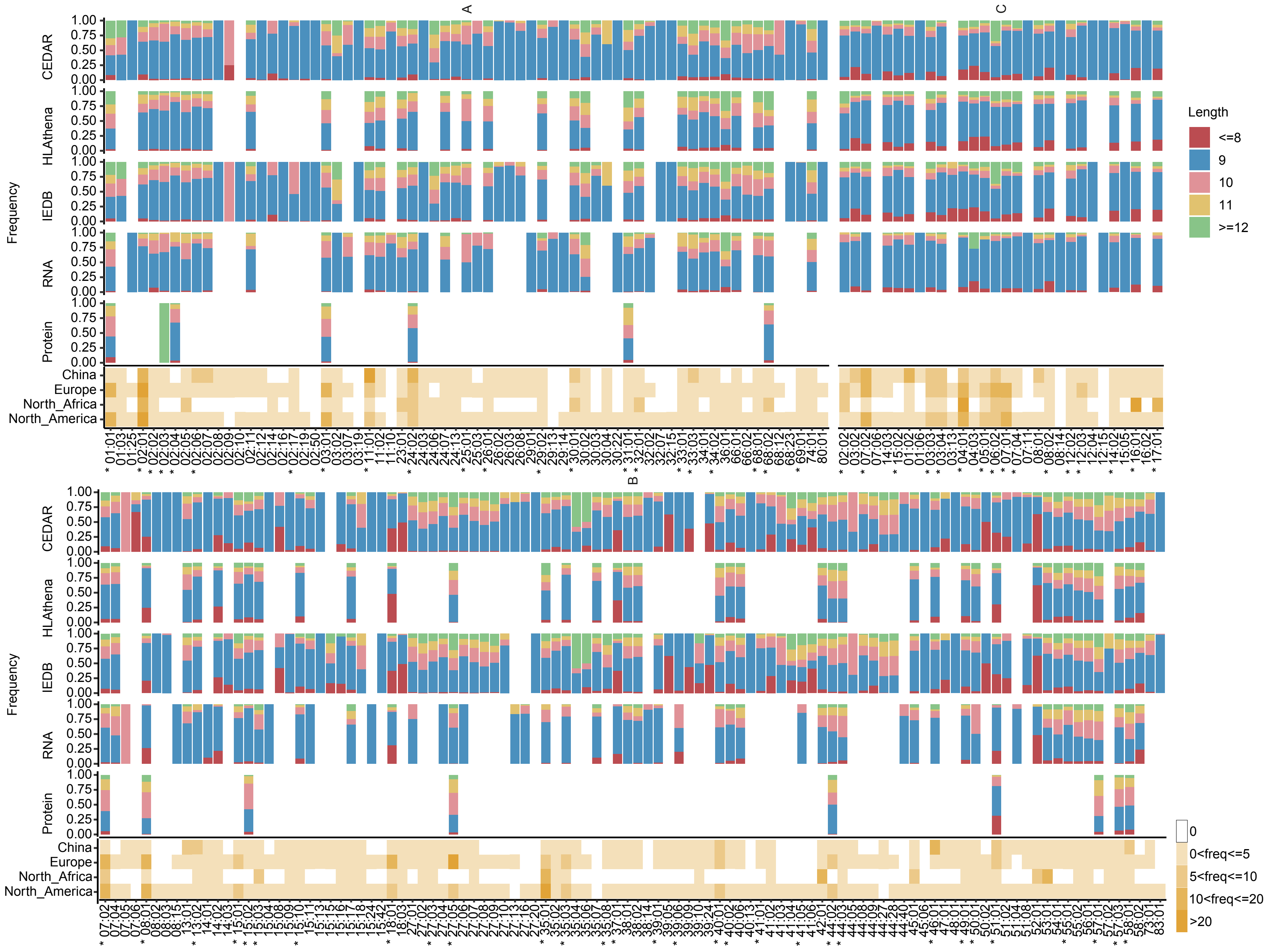

### Supplementary Figure 3

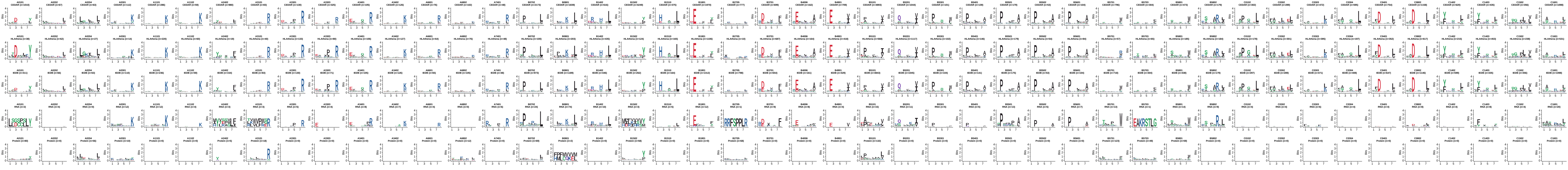

### Supplementary Figure 4

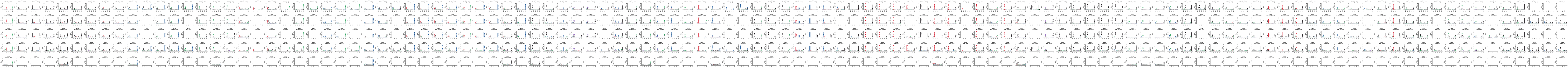

### Supplementary Figure 5

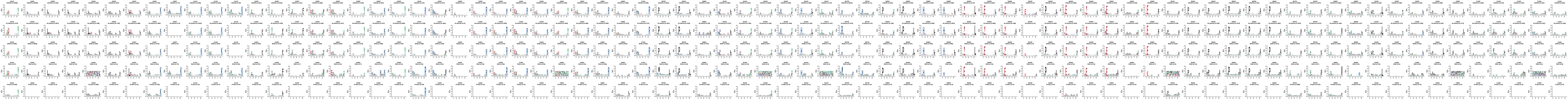

### Supplementary Figure 6

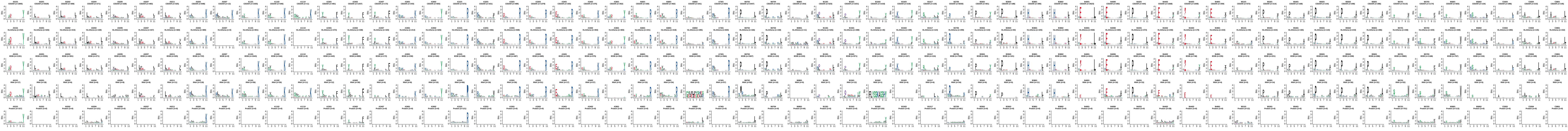

### Supplementary Figure 8

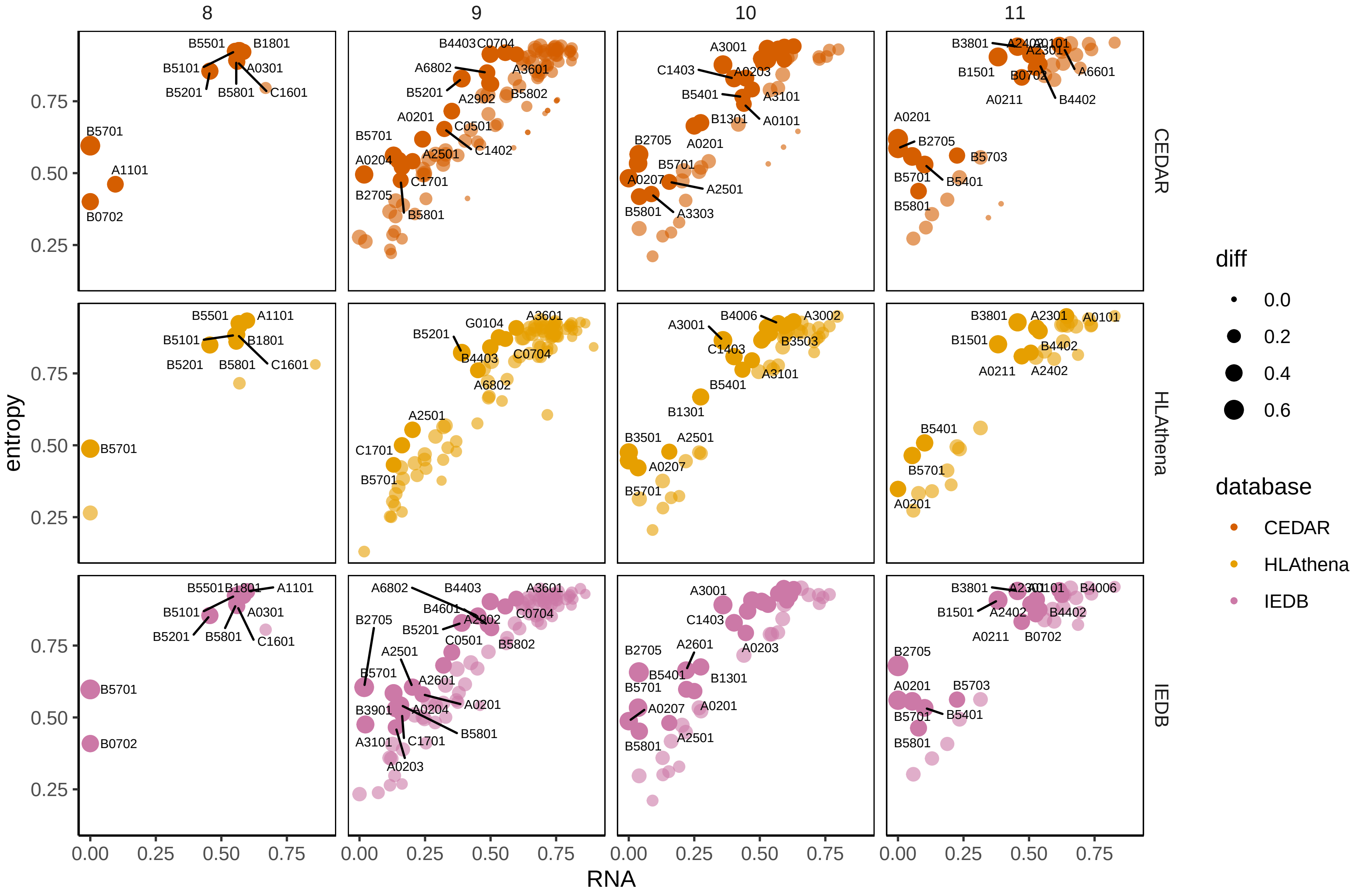

### Supplementary Figure 9

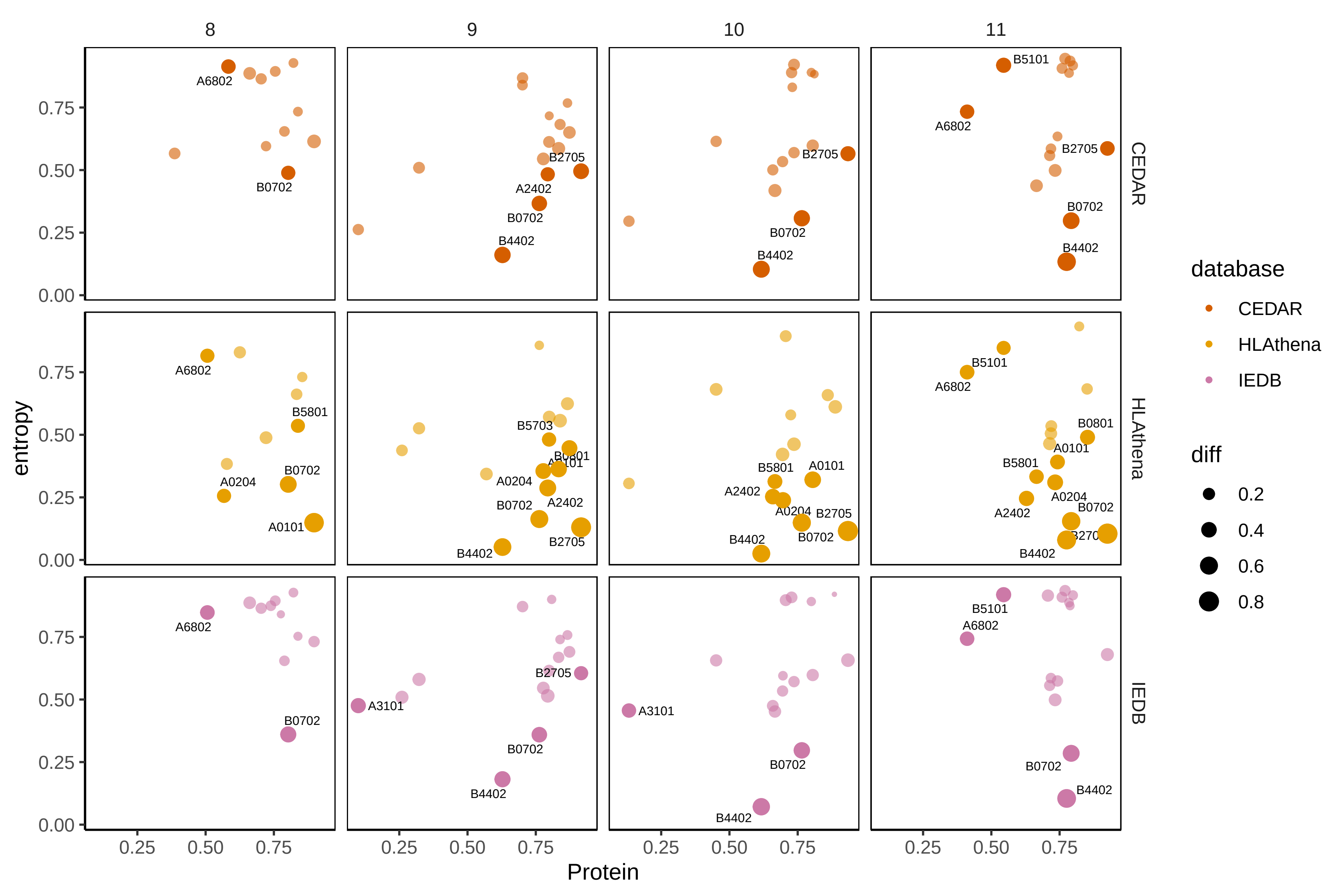

### Supplementary Figure 10

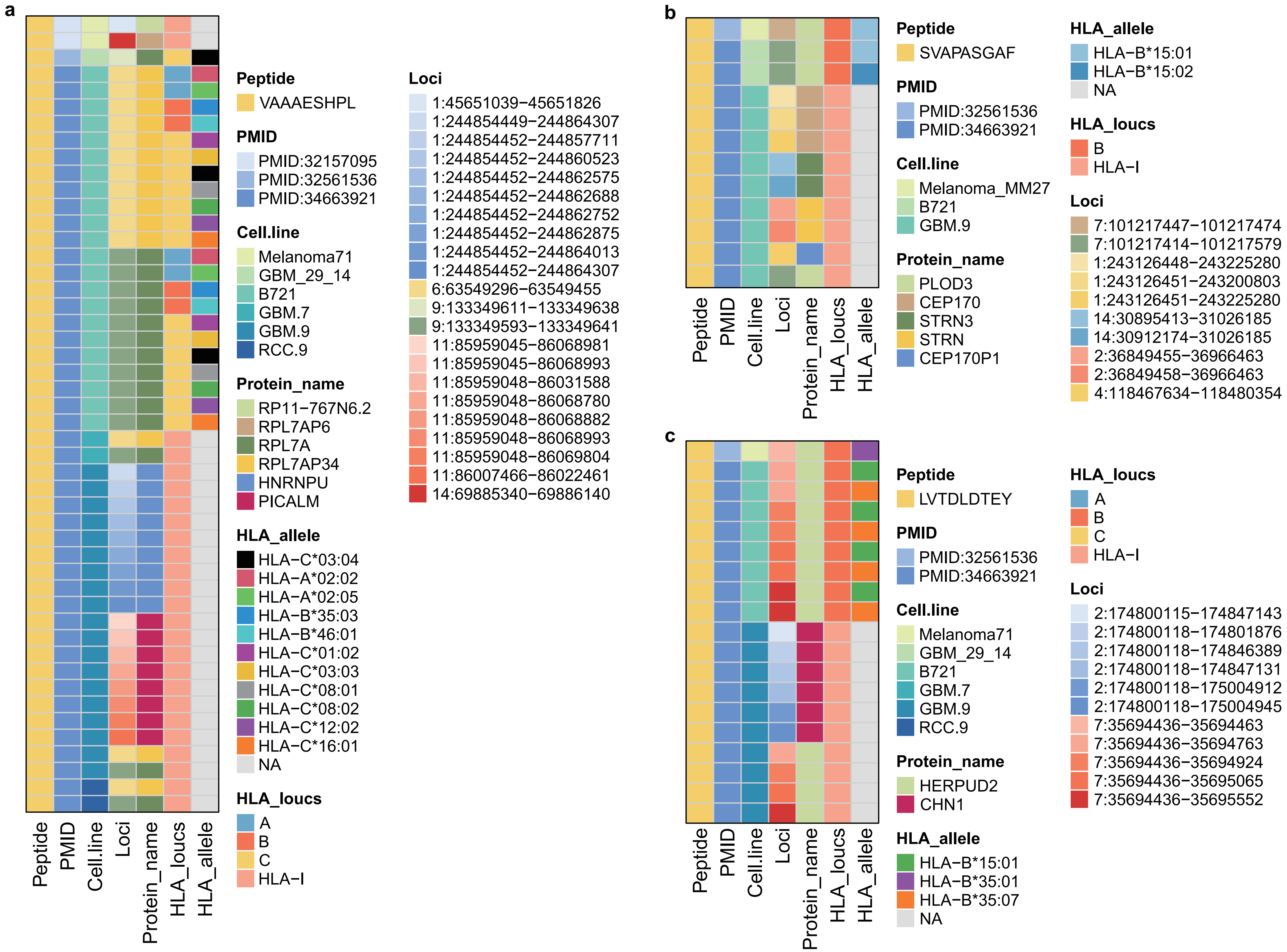
