## Supplementary Figure 7 for "NeoAtlas and NeoBert: A Database and A Predictive Model for Canonical and Noncanonical Tumor Neoantigens"

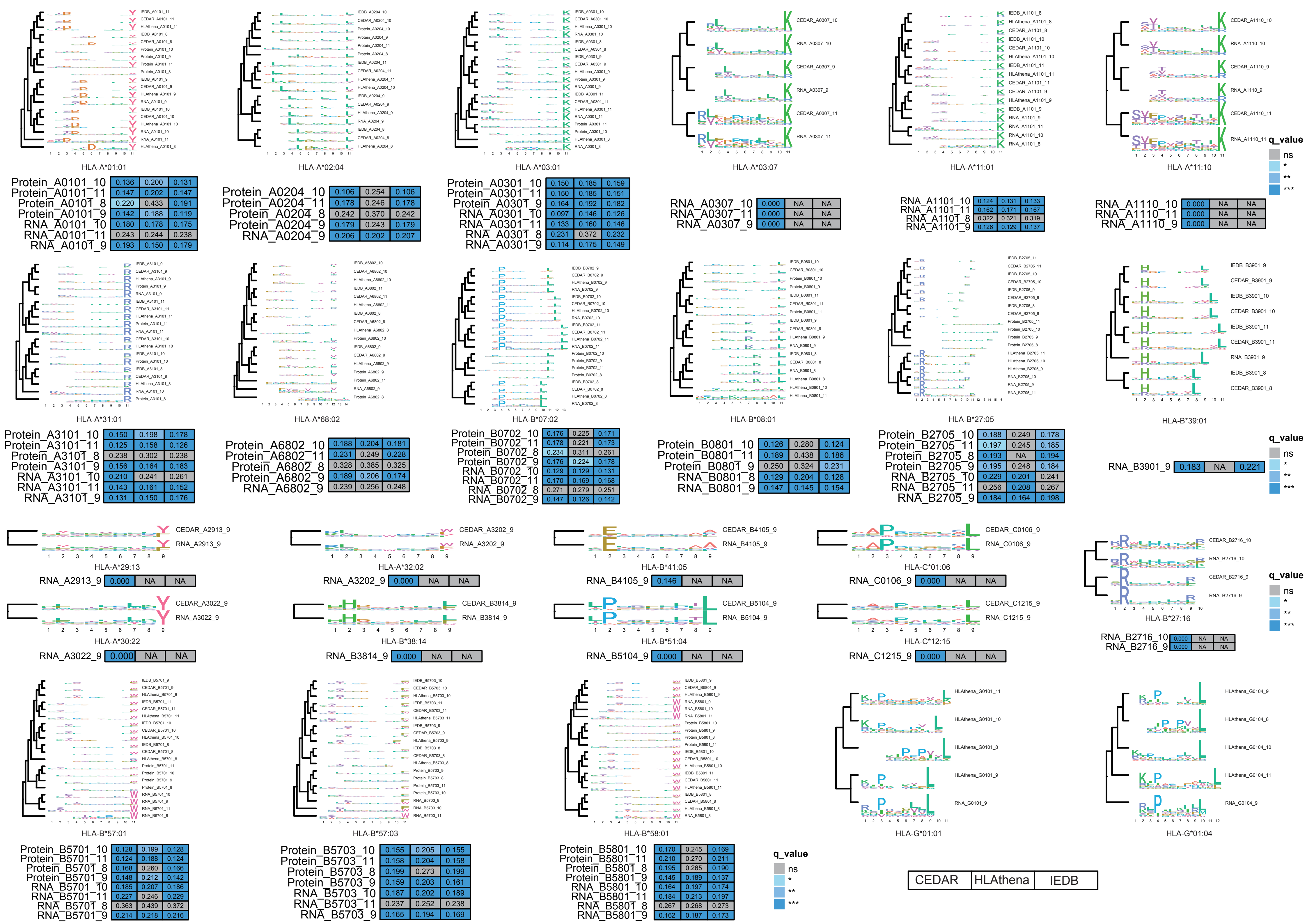


| | | | |
|---|---|---|---|
| Protein_A0101_10 | 0.136 | 0.200 | 0.131 |
| Protein_A0101_11 | 0.147 | 0.202 | 0.147 |
| Protein_A0101_8 | 0.220 | 0.433 | 0.191 |
| Protein_A0101_9 | 0.142 | 0.188 | 0.119 |
| RNA_A0101_10 | 0.180 | 0.178 | 0.175 |
| RNA_A0101_11 | 0.243 | 0.244 | 0.238 |
| RNA_A0101_9 | 0.193 | 0.150 | 0.179 |

| | | | |
|---|---|---|---|
| Protein_A0204_10 | 0.106 | 0.254 | 0.106 |
| Protein_A0204_11 | 0.178 | 0.246 | 0.178 |
| Protein_A0204_8 | 0.242 | 0.370 | 0.242 |
| Protein_A0204_9 | 0.179 | 0.243 | 0.179 |
| RNA_A0204_9 | 0.206 | 0.202 | 0.207 |

| | | | |
|---|---|---|---|
| Protein_A0301_10 | 0.150 | 0.185 | 0.159 |
| Protein_A0301_11 | 0.150 | 0.185 | 0.151 |
| Protein_A0301_9 | 0.164 | 0.192 | 0.182 |
| RNA_A0301_10 | 0.097 | 0.146 | 0.126 |
| RNA_A0301_11 | 0.133 | 0.160 | 0.146 |
| RNA_A0301_8 | 0.231 | 0.372 | 0.232 |
| RNA_A0301_9 | 0.114 | 0.175 | 0.149 |

| | | | |
|---|---|---|---|
| RNA_A0307_10 | 0.000 | NA | NA |
| RNA_A0307_11 | 0.000 | NA | NA |
| RNA_A0307_9 | 0.000 | NA | NA |

| | | | |
|---|---|---|---|
| RNA_A1101_10 | 0.124 | 0.131 | 0.133 |
| RNA_A1101_11 | 0.162 | 0.171 | 0.167 |
| RNA_A1101_8 | 0.322 | 0.321 | 0.319 |
| RNA_A1101_9 | 0.126 | 0.129 | 0.137 |

| | | | |
|---|---|---|---|
| RNA_A1110_10 | 0.000 | NA | NA |
| RNA_A1110_11 | 0.000 | NA | NA |
| RNA_A1110_9 | 0.000 | NA | NA |

| | | | |
|---|---|---|---|
| Protein_A3101_10 | 0.150 | 0.198 | 0.178 |
| Protein_A3101_11 | 0.125 | 0.158 | 0.126 |
| Protein_A3101_8 | 0.238 | 0.302 | 0.238 |
| Protein_A3101_9 | 0.156 | 0.164 | 0.183 |
| RNA_A3101_10 | 0.210 | 0.241 | 0.261 |
| RNA_A3101_11 | 0.143 | 0.161 | 0.152 |
| RNA_A3101_9 | 0.131 | 0.150 | 0.176 |

| | | | |
|---|---|---|---|
| Protein_A6802_10 | 0.188 | 0.204 | 0.181 |
| Protein_A6802_11 | 0.231 | 0.249 | 0.228 |
| Protein_A6802_8 | 0.328 | 0.385 | 0.325 |
| Protein_A6802_9 | 0.189 | 0.206 | 0.174 |
| RNA_A6802_9 | 0.239 | 0.256 | 0.248 |

| | | | |
|---|---|---|---|
| Protein_B0702_10 | 0.176 | 0.225 | 0.171 |
| Protein_B0702_11 | 0.178 | 0.221 | 0.173 |
| Protein_B0702_8 | 0.234 | 0.311 | 0.261 |
| Protein_B0702_9 | 0.176 | 0.224 | 0.178 |
| RNA_B0702_10 | 0.129 | 0.129 | 0.131 |
| RNA_B0702_11 | 0.170 | 0.169 | 0.168 |
| RNA_B0702_8 | 0.271 | 0.279 | 0.251 |
| RNA_B0702_9 | 0.147 | 0.126 | 0.142 |

| | | | |
|---|---|---|---|
| Protein_B0801_10 | 0.126 | 0.280 | 0.124 |
| Protein_B0801_11 | 0.189 | 0.438 | 0.186 |
| Protein_B0801_9 | 0.250 | 0.324 | 0.231 |
| RNA_B0801_8 | 0.129 | 0.204 | 0.128 |
| RNA_B0801_9 | 0.147 | 0.145 | 0.154 |

| | | | |
|---|---|---|---|
| Protein_B2705_10 | 0.188 | 0.249 | 0.178 |
| Protein_B2705_11 | 0.197 | 0.245 | 0.185 |
| Protein_B2705_8 | 0.193 | NA | 0.194 |
| Protein_B2705_9 | 0.195 | 0.248 | 0.184 |
| RNA_B2705_10 | 0.229 | 0.201 | 0.241 |
| RNA_B2705_11 | 0.256 | 0.208 | 0.267 |
| RNA_B2705_9 | 0.184 | 0.164 | 0.198 |

| | | | |
|---|---|---|---|
| RNA_B3901_9 | 0.183 | NA | 0.221 |

| | | | |
|---|---|---|---|
| RNA_A2913_9 | 0.000 | NA | NA |

| | | | |
|---|---|---|---|
| RNA_A3202_9 | 0.000 | NA | NA |

| | | | |
|---|---|---|---|
| RNA_B4105_9 | 0.146 | NA | NA |

| | | | |
|---|---|---|---|
| RNA_C0106_9 | 0.000 | NA | NA |

| | | | |
|---|---|---|---|
| RNA_B2716_10 | 0.000 | NA | NA |
| RNA_B2716_9 | 0.000 | NA | NA |

| | | | |
|---|---|---|---|
| RNA_A3022_9 | 0.000 | NA | NA |

| | | | |
|---|---|---|---|
| RNA_B3814_9 | 0.000 | NA | NA |

| | | | |
|---|---|---|---|
| RNA_B5104_9 | 0.000 | NA | NA |

| | | | |
|---|---|---|---|
| RNA_C1215_9 | 0.000 | NA | NA |

| | | | |
|---|---|---|---|
| Protein_B5701_10 | 0.128 | 0.199 | 0.128 |
| Protein_B5701_11 | 0.124 | 0.188 | 0.124 |
| Protein_B5701_8 | 0.168 | 0.260 | 0.166 |
| Protein_B5701_9 | 0.148 | 0.212 | 0.142 |
| RNA_B5701_10 | 0.185 | 0.207 | 0.186 |
| RNA_B5701_11 | 0.227 | 0.246 | 0.229 |
| RNA_B5701_8 | 0.363 | 0.439 | 0.372 |
| RNA_B5701_9 | 0.214 | 0.218 | 0.216 |

| | | | |
|---|---|---|---|
| Protein_B5703_10 | 0.155 | 0.205 | 0.155 |
| Protein_B5703_11 | 0.158 | 0.204 | 0.158 |
| Protein_B5703_8 | 0.199 | 0.273 | 0.199 |
| Protein_B5703_9 | 0.159 | 0.203 | 0.161 |
| RNA_B5703_10 | 0.187 | 0.202 | 0.189 |
| RNA_B5703_11 | 0.237 | 0.252 | 0.238 |
| RNA_B5703_9 | 0.165 | 0.194 | 0.169 |

| | | | |
|---|---|---|---|
| Protein_B5801_10 | 0.170 | 0.245 | 0.169 |
| Protein_B5801_11 | 0.210 | 0.270 | 0.211 |
| Protein_B5801_8 | 0.195 | 0.265 | 0.190 |
| Protein_B5801_9 | 0.145 | 0.189 | 0.137 |
| RNA_B5801_10 | 0.164 | 0.197 | 0.174 |
| RNA_B5801_11 | 0.184 | 0.213 | 0.197 |
| RNA_B5801_8 | 0.267 | 0.268 | 0.273 |
| RNA_B5801_9 | 0.162 | 0.187 | 0.173 |
